# Expression and Function of Connexin 43 and Connexin 37 in the Murine Zona Glomerulosa

**DOI:** 10.1101/2024.08.26.608761

**Authors:** Gabriel Stölting, Nicole Hellmig, Hoang An Dinh, Marina Volkert, Ute I. Scholl

**Affiliations:** Berlin Institute of Health at Charité – Universitätsmedizin Berlin, Center of Functional Genomics, Charitéplatz 1, 10117 Berlin, Germany; Charité – Universitätsmedizin Berlin, Institute of Translational Physiology, Charitéplatz 1, 10117 Berlin, Germany

**Keywords:** aldosterone, gap junction, connexin, zona glomerulosa, calcium imaging

## Abstract

The zona glomerulosa (ZG) synthesizes the mineralocorticoid aldosterone. The primary role of aldosterone is the maintenance of volume and electrolyte homeostasis. Aldosterone synthesis is primarily regulated via tightly controlled oscillations in intracellular calcium levels in response to stimulation. It has previously been shown that calcium oscillations are synchronized through mechanical linkage between adjacent ZG cells. In many other cell types, similar synchronization is rather dependent on gap junctions (GJ). The recent discovery of mutations in *CADM1* was linked to impaired GJ function in the ZG. Based on published transcriptomics data, we re-examined the presence and functional impact of GJ in the ZG. We found evidence for the expression of connexin 43 and 37 in the ZG in microarray data, in-situ hybridization and immunohistology. Calcium oscillations in ZG rosettes showed some degree of synchronization as reported previously. Unspecific GJ inhibition only had a small impact on this synchronicity. However, no signs of connections between cytosols could be observed as indicated by the lack of fluorescence recovery after photobleaching. We conclude that, while connexin proteins are expressed in the ZG, functional GJ in the physiological ZG are rare and of little consequence for calcium signaling.

## INTRODUCTION

The function of the adrenal cortex is the synthesis of steroid hormones, ranging from androgens in the zona reticularis (ZR; not present in mice) and cortisol (or corticosterone in mice) in the zona fasciculata (ZF) to aldosterone in the zona glomerulosa (ZG). As early as fifty years ago, an extensive network of gap junctions was observed in the adrenal cortex (Friend and Gilula, 1972). These protein complexes form pores between adjacent cells that permit the transfer of ions and larger molecules as well as electrical signals across cellular barriers. Molecularly, they are formed by the assembly of two hemichannels (connexons), one from each of the adjacent cell membranes. Each connexon is a hexamer of proteins from the large and widely expressed connexin family (Kumar and Gilula, 1996).

Previous studies have mostly implicated connexin 43 (Cx43, Gene: *Gja1*) as the central isoform in the formation of gap junctions across the adrenal cortex (Murray et al., 1995; Murray and Pharrams, 1997). Coupling was reported to be abundant in the ZF and ZR (Murray et al., 1995; Davis et al., 2002). Within the ZF, coupling via gap junctions was suggested to increase sensitivity to ACTH via transfer of cAMP between cells (Munari-Silem et al., 1995, p.) but also to attenuate cellular proliferation (Bell and Murray, 2016).

Several previous studies have shown that Cx43 is only expressed at very low levels in the ZG of most species, if at all (Murray et al., 1995; Murray and Pharrams, 1997; Davis et al., 2002) while other studies suggest relevant expression in the ZG (Meda et al., 1993). The investigation of the expression of other connexin isoforms is less complete (for a review see (Bell and Murray, 2016)). These results have led to the assumption that gap junctions do not play a major role in the function of the ZG. Recent studies, however, have demonstrated coupling of calcium signals across adjacent ZG cells (Guagliardo et al., 2020), and new studies of the human ZG transcriptome have suggested expression of connexin isoform Cx43, but also connexin 37 (Cx37, Gene: *Gja4*) (Nishimoto et al., 2012, 2015).

Further interest in this question came from the recent discovery of primary aldosteronism-associated mutations in *CADM1* (Wu et al., 2023). The encoded protein was postulated to play a role in establishing cell-cell contacts in other tissues (Ito et al., 2012), and the discovered mutations are suggested to cause a change in the angle of these contacts, placing cells further apart. This was suggested to lead to the inability to form gap junctions, underlying the increased synthesis of aldosterone in the ZG (Wu et al., 2023).

Calcium is the most important messenger in ZG cells, regulating the expression of enzymes involved in aldosterone synthesis as well as in the transport of cholesterol into mitochondria (Fakunding et al., 1979; Fern et al., 1995; Pezzi et al., 1997; Bassett et al., 2004). Others and we have demonstrated that the intracellular calcium concentration in the ZG is regulated by modulating the oscillatory influx of calcium via electrical depolarizations. This increase is mediated by calcium spikes with a frequency of around 1 Hz that are typically clustered into bursts of activity (Schewe et al., 2019; Guagliardo et al., 2020; Dinh et al., 2024). This is predominantly seen in slice preparations as dissociated cells seem to lose the propensity for calcium spiking even though many of the important membrane conductances are retained (Hu et al., 2012; Penton et al., 2012). Calcium influx itself seems to be primarily mediated by voltage-gated calcium channels which are also indispensable for spiking (Hu et al., 2012; Stölting et al., 2023; Dinh et al., 2024). The magnitude of the two main stimuli of aldosterone synthesis, the serum concentrations of angiotensin II and potassium, are transduced into increases of the intracellular calcium concentration of the ZG (Capponi et al., 1984) via increases in spiking (Schewe et al., 2019; Guagliardo et al., 2020). As the name suggests, cells within the ZG are arranged in spherical arrangements of several cells; these clusters are enclosed by laminin-rich basal membranes that separate glomeruli. A temporal correlation between the oscillation pattern between ZG cells has recently been demonstrated and analyzed, but was attributed to a mechanical linkage through beta-catenin rather than electrical or chemical coupling via gap junction proteins, based on the previously reported lack of gap junctions in the ZG (Guagliardo et al., 2020). Given this data and interest in the maintenance of electrical and calcium signals within cells of the ZG in humans but also murine model systems as well as the proposed link between *CADM1* mutations and gap junction formation, we here re-investigate the presence and potential role of gap junctions in the murine ZG using current histology and imaging techniques.

## MATERIALS & METHODS

### Animals

All mice used were kept under specific pathogen-free conditions at the Forschungseinrichtungen für Experimentelle Medizin (Charité – Universitätsmedizin Berlin) according to all relevant regulations. This includes a 12h light/dark cycle with free access to food and water as well as enrichment materials.

### Analysis of microarray data

Microarray analysis data was obtained from the supplementary information in Nishimoto et al. (Nishimoto et al., 2015). Data was loaded via a custom python script, and expression values were taken from the “ZG” and “ZF” columns of the original excel file. Data was log10 transformed for visualization, and genes were identified via their gene names.

### RNA in-situ hybridization

In situ hybridization was performed on 5 µm sections of formalin-fixed, paraffin-embedded adrenal glands using the RNAscope 2.5 HD Assay – Brown (Advanced Cell Diagnostics) according to the instructions. Probes used were: Mm-Gja1 (Cat No. 486191), Mm-Gja4 (Cat No. 588591), Mm-Ppib (positive control, Cat No. 313911) and DapB (negative control, Cat No. 310043). Images were taken on a Zeiss Axioplan 2 microscope with a 20x/0.45 or 40x/0.75 objective.

### Immunofluorescence

Mice were euthanized by cervical dislocation under isoflurane anesthesia as described above. For some mice, postmortem perfusion, first with saline later with formalin, was initiated immediately after euthanasia. Afterwards, the adrenal glands were extracted and stored in formalin at 4 °C for 24 h. Following this period, formalin-fixed adrenal glands were dehydrated in increasing concentrations of ethanol (70%, 80%, 90%, 95% and 100%) for 60 minutes each. For paraffin embedding, samples were transferred 2x to 100% xylene before embedding in paraffin resulting in formalin-fixed paraffin-embedded (FFPE) tissue samples. FFPE blocks were cut on 5 µm thick slices on a microtome (RM2125, Leica Instruments) and mounted onto microscopy slides (SuperFrost Plus, Menzel Gläser). Before staining, slides were deparaffinized by immersion in xylene for 5 minutes, followed by rehydration in decreasing concentrations of ethanol (100%, 95%, 90%, 80% and 70%) and finally deionized water for 2-5 minutes each. For the Cx37 peptide control, samples were incubated before staining (Aviva, ARP36603, 1 mg/ml). Samples were incubated with primary antibodies (Connexin 43: C6219, Sigma-Aldrich, dilution 1:500; Connexin 37: ARP36603-P050, Aviva, dilution 1:300; Dab2: sc-136964, Santa Cruz, dilution: 1:400) overnight, followed by washing 3x with TBS and incubation with the secondary antibodies (Anti-mouse Alexa fluor 488: A21202, Invitrogen, dilution 1:400; Anti-rabbit Alexa fluor 647: A32728, Invitrogen, dilution 1:400) for 1 h at room temperature (∼21 °C). All samples were mounted in Vectashield with DAPI (Vector Laboratories) and imaged on a Zeiss Axio Observer widefield flurorescence microscope using a 20x, NA 0.8 objective.

### Calcium imaging in acute adrenal slice preparations

Acute adrenal slices were prepared as described previously (Schewe et al., 2019; Seidel et al., 2021). Murine (C57BL/6N) adrenal glands from 12-20 week-old mice were rapidly extracted following cervical dislocation under isoflurane anesthesia and transferred to bicarbonate-buffered saline (BBS in mM: 100 NaCl, 26 NaHCO_3_, 10 D-Glucose, 10 HEPES, 5MgCl_2_, 2 KCl) on ice. Glands were embedded in 3% low-melting temperature agarose dissolved in BBS prep and mounted on a vibratome (7000 smz-2; Campden Instruments). Afterwards, slices were maintained in BBS supplemented with 2 mM CaCl_2_. All solutions were continuously gassed with carbogen (95% O_2_, 5% CO_2_).

Staining of slices was performed for 1 hour in a cell culture insert with 750 µl of BBS on the outside and initially 64 µM Fura-2 + 10% Pluronic F-127 dissolved in BBS on the inside. Recordings were performed on a SliceScope (Scientifica) with a CoolLED light source (Cairn Research), and images were taken every 100 ms with an exposure of 10 ms using an OptiMOS camera (Qimaging) controlled by MicroManager (https://micro-manager.org). Recording solution was BBS containing a total of 4 mM K^+^. The first minute following a solution change was omitted from analysis to allow for complete solution exchange, even deeper in the tissue.

### Fluorescence recovery after photobleaching

For fluorescence recovery after photobleaching experiments, acute slice preparations were prepared as described above. Staining was performed for 1 hour using 1µM Calcein AM. Images were taken on a Nikon spinning disk confocal CSU-W1 SoRa using a 40x, NA 1.25 objective with silicone immersion. Bleaching and imaging was performed using the 488 nm laser line and recorded using a Hamamatsu ORCA-Fusion camera. For some experiments, calcium signals were simultaneously recorded to assess slice viability using Calbryte 630 (AAT Bioquest). Staining was performed by adding 37 µM Calbryte + 0.001% Pluronic F-127 to the Calcein AM staining solution. Recording was performed using the 594 nm laser line and a second Hamamatsu ORCA-Fusion camera.

### Analysis of calcium imaging

Recordings were loaded into Fiji (Schindelin et al., 2012) and drift corrected using the “manual drift correction” plugin. Cells were manually selected, and intensity profiles over time extracted for further analysis. Data was loaded into a custom written software, and calcium spikes were automatically detected under manual supervision. Further analyses were performed using custom-written python scripts. Temporal similarity of spike trains was taken as a measure of synchronized activity. To quantify this behavior, we utilized the Jaccard index (from SciPy (Virtanen et al., 2020)), which is defined as the ratio of the number of temporally synchronized spikes by the number of spikes overall across two cells. Confidence intervals were calculated by bootstrap resampling with 10,000 resamples unless specified otherwise.

## RESULTS

### Presence of connexin mRNA in the murine ZG

We re-analyzed publicly available microarray data from micro-dissected human adrenal cortex (Nishimoto et al., 2015). Two connexin isoforms, *GJA1* (encoding connexin 43) and *GJA4* (encoding connexin 37; Figure 1A), showed the highest expression. *GJA1* was more strongly expressed than *GJA4* in both, ZG and ZF. Whereas *GJA1* expression increased towards the ZF, *GJA4* showed slightly higher expression in the ZG than in ZF (Figure 1A). This is also supported by transcriptome data of the rat adrenal cortex from the same group (Nishimoto et al., 2012). To assess whether the signal seen in micro-dissected tissue arises from adrenocortical cells or from blood vessels of the adrenal cortex (Lai et al., 2020), we performed in-situ hybridization (ISH), which provides high spatial resolution and sensitivity to associate the presence of mRNAs of interest to specific cells. ISH in formalin-fixed, paraffin-embedded slices from mouse adrenal glands confirmed the occurrence of mRNA for *Gja1* (Cx43) in the adrenal cortex (Figure 1B left). *Gja1* signals increased towards the interior of the adrenal gland, confirming earlier reports of Cx43 being predominantly expressed in the murine ZF. However, some signals could also be seen in cells of the ZG, and strong signals were present in the capsular region (Figure 1B). Based on the microarray data, we also examined the presence of *Gja4* in the adrenal cortex (Figure 1B right). Signals could be seen in endothelial cells delineating small blood vessels of the adrenal cortex (asterisk in Figure 1B right), which have previously been described (Lai et al., 2020), but also in glomerulosa cells. Some *Gja4* transcripts appeared in cells of the ZF, particularly in cells closer to the ZG. In contrast to *Gja1*, we found the expression of *Gja4* to be lower in the capsular region. Positive and negative controls are shown in Supplementary Figure S1.

**FIGURE 1.**
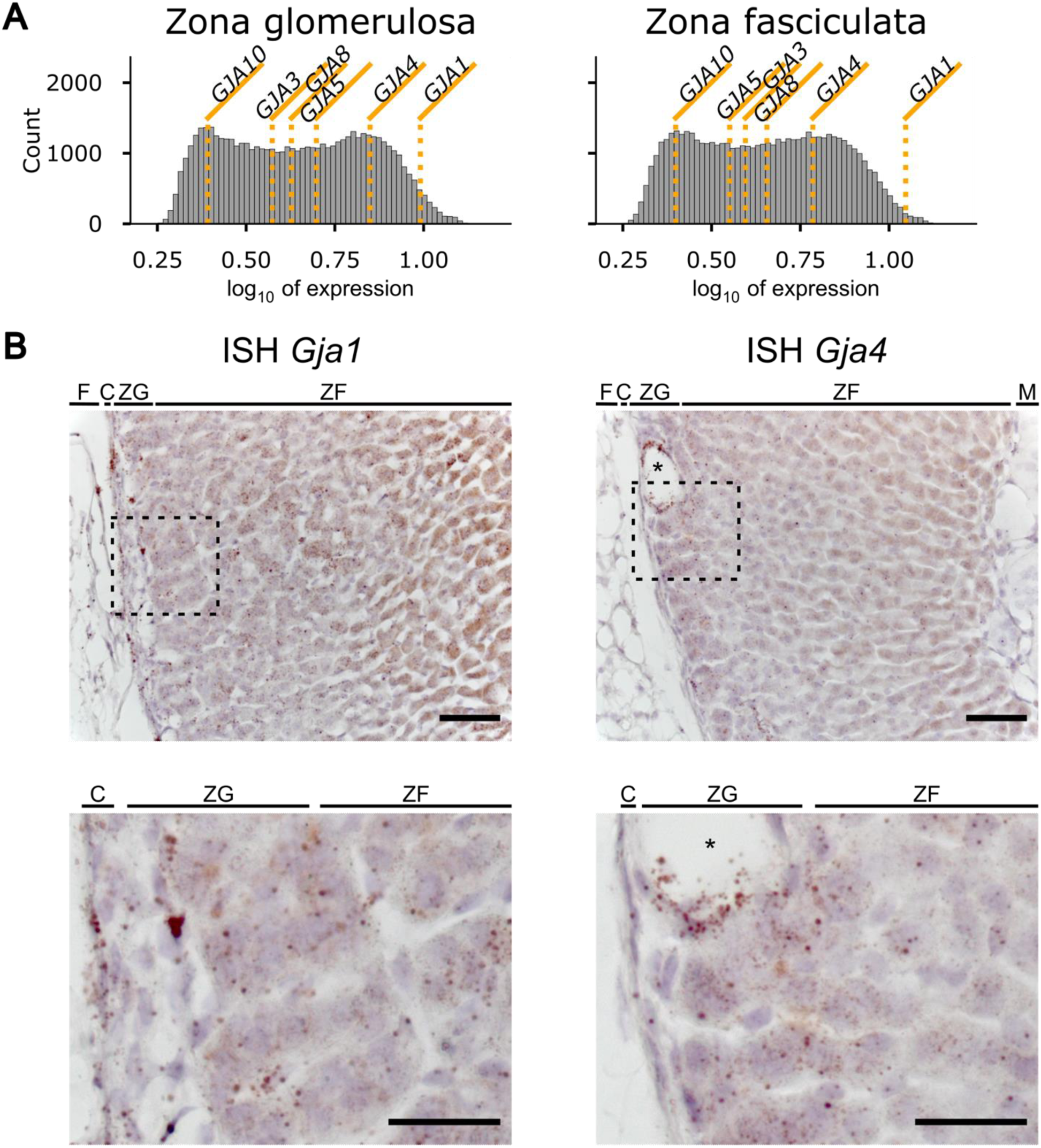
*Gja1* (connexin 43) and *Gja4* (connexin 37) mRNA is present in the adrenal zona glomerulosa. **(A)** Analysis of the microarray analysis of human adrenal cortices by (Nishimoto et al., 2015). Mean expression values for various connexin isoforms are plotted over a histogram of the expression values for all determined genes within the array. *Gja1* (connexin 43) and *Gja4* (connexin 37) are the most abundantly expressed connexin isoforms. **(B)** ISH revealed mRNA of both *Gja1* and *Gja4* in the ZG and ZF of the adrenal cortex. Brown spots indicating mRNA for *Gja1* can be seen prominently in fibroblasts of the capsule as well as in adrenocortical cells, increasing in density towards the medulla. *Gja4* mRNA could not be observed in the capsule but prominently in the ZG and in cells delineating blood vessels of the adrenal cortex (*). Dashed boxes indicate magnified regions shown at the bottom. Brown spots represent signals from stained mRNA molecules. Images are representative of ISH experiments on slices from three individual animals each (2 male, 1 female). Scale bar, 50 µm or 25 µm for the magnified image. F – adipose tissue, C – capsule, ZG – zona glomerulosa, ZF – zona fasciculata, M – medulla.

### Presence of connexin proteins in the murine ZG

We used immunofluorescence (IF) staining to investigate the expression of Cx43 and Cx37 in the ZG. For the staining of connexin 43, we used an antibody that was previously used in several studies investigating conditional knock-out (KO) models (Clasadonte et al., 2017; Christopher et al., 2022). Overall, the signal was strongest in the adrenal medulla, almost completely masking the signal in the cortex (Figure 2A). Adjusting the signal range to the intensities observed in the adrenal cortex led to observations that agree with previous studies (Murray and Pharrams, 1997; Davis et al., 2002; Wu et al., 2023): The Cx43 signal in the ZG was present but lower than in the ZF (Figure 2B) and clearly different than in controls using secondary antibody only (Supplementary Figure S2A, B).

**FIGURE 2.**
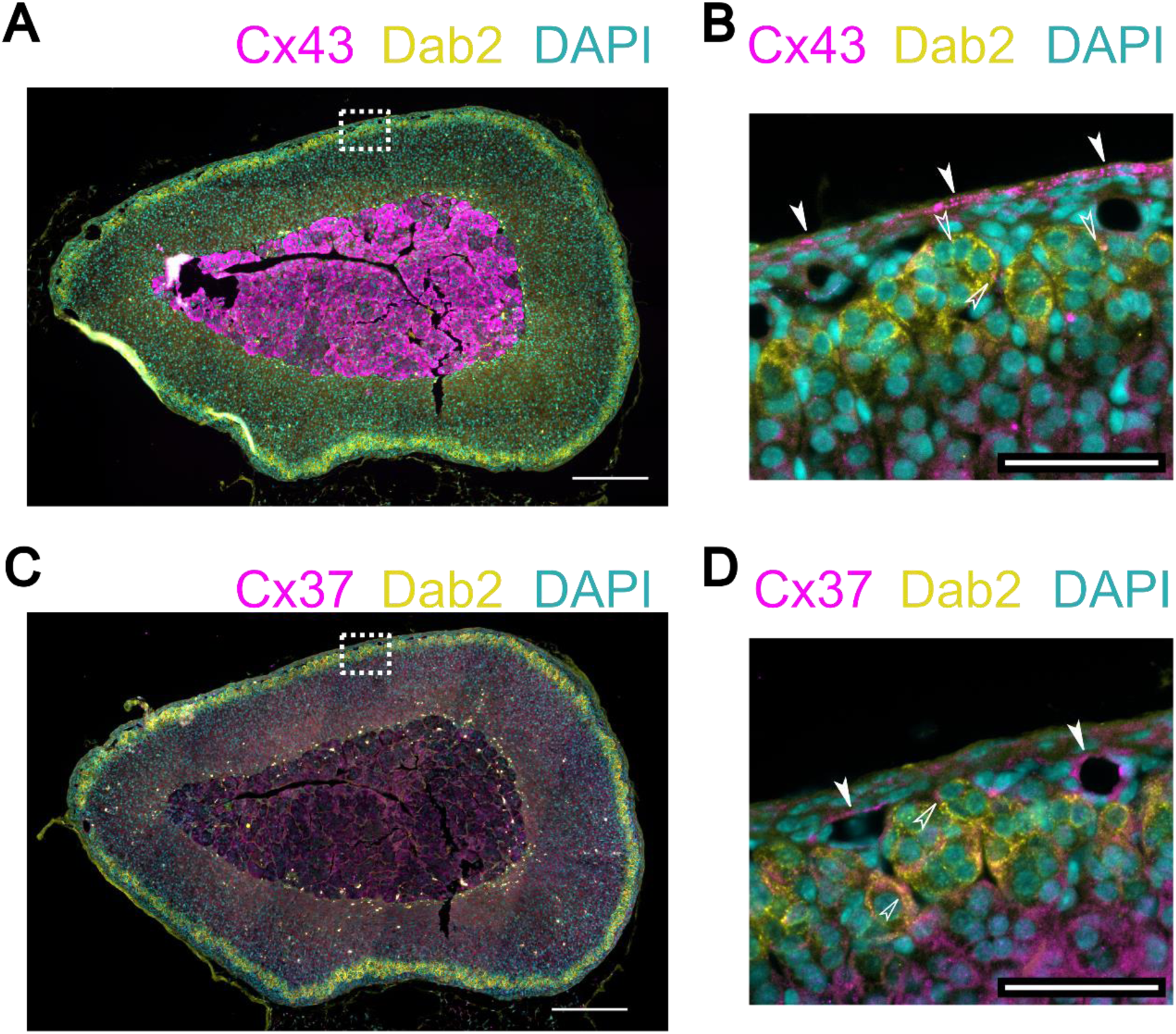
Connexin 43 and 37 proteins are present in the adrenal cortex. **(A)** Immunofluorescence staining reveals connexin 43 (magenta) predominantly in the medulla. Smaller signals are observed in the zona fasciculata (ZF) and even less in the zona glomerulosa (ZG; defined by staining of Dab2 (yellow)). Nuclei are stained with DAPI (cyan). The image is representative of the staining of sections (5 µm thickness) from 8 animals. Scale bar: 250 µm. **(B)** Magnification of the part enclosed by the white dashed square in A. Cx43 signals are predominantly seen in the capsule (filled arrows) but also in the ZG (open arrows) and further into the ZF at the bottom. Scale bar: 50 µm. **(C)** Immunofluorescence staining reveals low expression of connexin 37 overall (magenta). The zona glomerulosa (ZG) is again highlighted by staining of Dab2 (yellow). Nuclei are stained with DAPI (cyan). The image is representative of the staining of sections (5 µm thickness) from 8 animals. Scale bar: 250 µm. **(D)** Magnification of the part enclosed by the white dashed square in A. Cx37 signals are seen around blood vessels (filled arrows) but also sparsely in the ZG (open arrows). Scale bar: 50 µm.

Staining for Cx37 also closely followed the distribution pattern observed in the ISH, with the staining intensity being highest around blood vessels with some staining in the adrenal cortex decreasing towards the medulla. (Figure 2C, D). Unlike for the chosen Cx43 antibody, this antibody has not been previously confirmed in KO samples. We performed a staining with an antigen peptide control, which greatly reduced overall fluorescence intensity and led to a different staining pattern (Supplementary Figure S2C, D). Our IF stainings suggest the presence of small amounts of both, Cx43 and Cx37 in the ZG but cannot provide proof of their functional relevance.

### Functional coupling of calcium signals

Stimulation of the ZG by angiotensin II or potassium is transduced into oscillations of intracellular calcium. The presence of gap junctions in the ZG may lead to a synchronization of these calcium signals between adjacent cells. Consistent with prior reports (Guagliardo et al., 2020), we observed some level of synchronized calcium spiking in neighboring cells (Figure 3A). We quantified the degree of synchronicity between spiking of different cells by comparing spike trains of all ZG cells observed within one experiment using the Jaccard index (JI; Figure 3B). The JI is a similarity coefficient ranging from 1.0 for perfectly identical spike trains to 0.0 for completely divergent activity patterns. Cells in the ZG are irregular in shape but mostly resemble rectangular to oval structures. To obtain their position for the calculation of distances between cells, we therefore fit a rectangle around each cell, the center of which was taken as center for the corresponding cell. While several cells showed synchronization, most spikes seemed to occur independently, and JIs between cells across the whole field of view were typically below 0.1 (Figure 3B and C). Higher synchronicity was limited to cells in close proximity (distance: 5.6 µm ± 0.8, mean ± s.e.m., n = 2413 calculated correlations, 186 cells, 8 slices, 6 animals; Figure 3C), very close to the 7.2 µm reported previously for coupled cells (Guagliardo et al., 2020), suggesting that coupling of signals mostly occurs among adjacent cells within ZG glomeruli.

**FIGURE 3.**
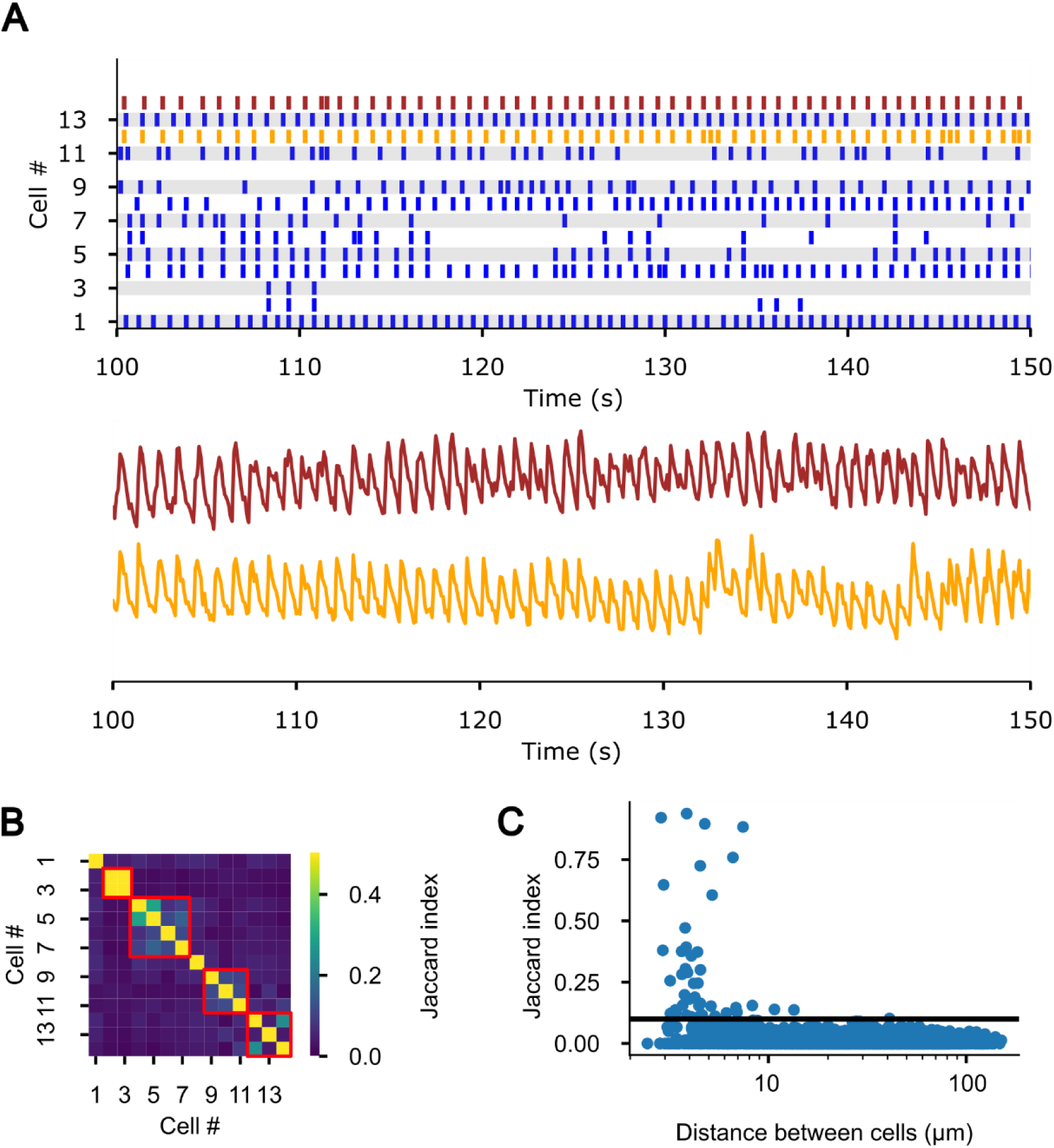
Calcium spiking is correlated in some cells of the murine ZG. **(A)** Representative spike trains over 500 frames (50 s at 10 frames/s) from 14 cells in an adrenal slice stained with the calcium indicator Calbryte 520 AM. Raw data from the spikes marked in yellow and red are shown below to highlight the synchronicity in spiking between these two cells. **(B)** Representative plot of the calculated Jaccard indices (JI) between all 14 cells in the same recording as shown in A. The full duration included in the analysis was 6600 frames (11 minutes). JI values are color coded as shown in the legend on the right. **(C)** JI from 186 cells (8 slices, 6 animals) are plotted against the distance (center to center) between the corresponding cells. The black line indicates the chosen cutoff of 0.1 for further analysis of cells with higher correlations.

However, only 66 out of 186 analyzed cells (8 slices, 6 animals) showed synchronicity with at least one other cell as defined by having at least one connection with a JI larger than 0.1. The mean JI for connections above this threshold was 0.36 (95% CI: 0.28-0.42, bootstrap resampling).

Spiking in ZG cells is highly regular, and correlation may therefore also arise by chance due to the finite camera frame rate that results in a binning effect of the raw calcium signal. To exclude random correlations, we calculated the JI with the same recording shifted by *i* frames (for *i* between 1 and 10) (Supplementary Figure S3). If the observed correlations were the result of random synchronicity, similar JI distributions would be observed in the shifted recordings. However, little to no synchronization could be observed in this analysis, supporting the veracity of the observed correlation in the previous analyses.

### Fluorescent signals in the ZG do not recover after photobleaching

Synchronization has previously been attributed to mechanical contacts dependent on cadherins (Guagliardo et al., 2020). Synchronization of calcium signals between cells thus does not serve as a proof of the presence of functional gap junctions. We used fluorescence recovery after photobleaching (FRAP) to study the presence of physical connections between ZG cells. In FRAP, adrenal slices were loaded with calcein AM (Figure 4A). Afterwards, the fluorescence of individual cells within the ZG was bleached using targeted high intensity laser illumination (Figure 4B). If cells were connected by gap junctions, non-bleached dye from adjacent cells should have flowed into the bleached cell, restoring the signal (Wade et al., 1986).

**FIGURE 4.**
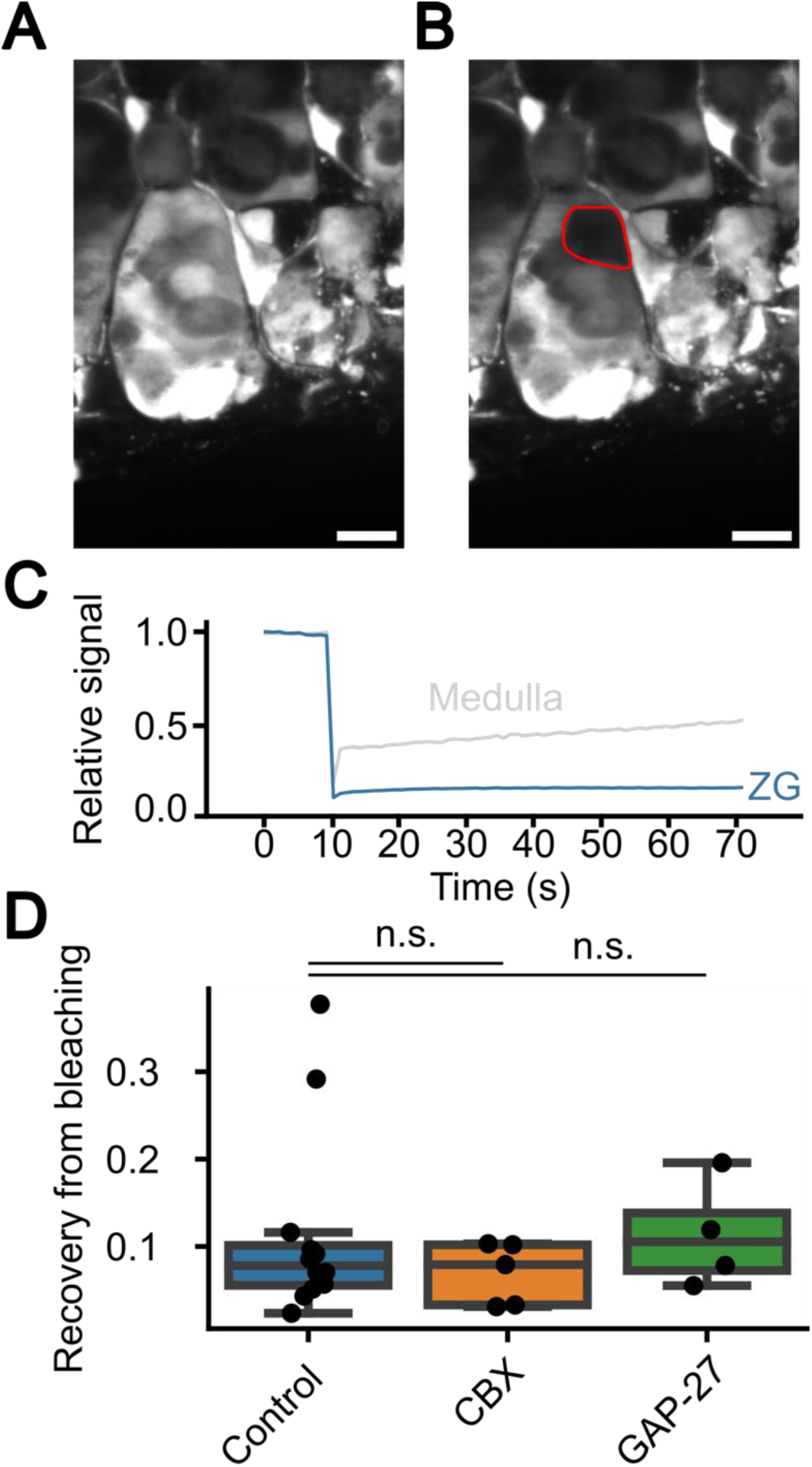
Murine zona glomerulosa cells show only minor cytosolic connections. **(A)** Representative image of Calcein AM-stained ZG cells within an acutely prepared murine adrenal gland slice before photobleaching. Scale bar: 10 µm. **(B)** The same field of view as in A after bleaching using 488 nm laser illumination. **(C)** Representative trace of a fluorescence recovery in the ZG (blue) and the adrenal medulla (grey) after a photobleaching experiment. The cells were stained with Calcein AM, and the fluorescence was bleached after 10 s using strong 488 nm laser illumination. The signal drops to approximately 10%, and recovery was observed over a minute. **(D)** Quantification of multiple FRAP experiments (Control; n = 12; mean = 0.11, sd = 0.10) reveals only low levels of fluorescence intensity recovery. Values are indistinguishable (n.s. – not significant) in the presence of the gap junction inhibitors carbenoxolone (CBX; n = 5; mean = 0.07, sd = 0.03; p = 0.38, t = 0.90, Student’s t-test) or GAP-27 (n=4; mean = 0.11, sd = 0.05; p = 0.97, t = 0.04, Student’s t-test), suggesting no involvement of gap junctions in the small recovery seen.

In general, we observed a recovery of only about 10% of the original signal in ZG cells (Figure 4C). Higher recovery was observed in the adrenal medulla (Figure 4C) which is known to contain an extensive network of gap junctions between chromaffin cells (Martin et al., 2001).

We also incubated slices with the known unspecific gap junction inhibitors carbenoxolone (CBX) (Davidson et al., 1986) or the more specific inhibitor GAP-27 (Chaytor et al., 1998) with no difference in recovery of the signal in ZG cells (Figure 4D).

Our results suggest that cells in the murine ZG are not connected by gap junctions under the conditions used in our experiments.

### Carbenoxolone only mildly reduces signal correlation

We also tested whether the observed temporal correlation of calcium signals decreased during perfusion with CBX. Application of the more specific blocker GAP-27 was not possible due to the amount of the substance that would be required for the chosen recording duration with constant perfusion at high flow rates (2-4 ml/min).

Application of 100 µM CBX via the extracellular perfusate reduced spiking in all cells (Figure 5A). Mean overall activity, as defined by the number of spikes per second, was reduced from 0.43 to 0.06 (Figure 5B). Spiking only slowly increased after washout on a similar time scale as also observed in slice preparations of other organs (Meme et al., 2009). Overall reduced spiking was due to complete cessation of spiking in some cells and reduced spiking and bursting in others (Figure 5A). For those cell-cell correlations with synchronization before application of CBX (JI > 0.1; Figure 5C), we observed a small reduction of the JI by 15.6% (Figure 5D). Almost no reduction of the JI was observed in controls without CBX (Figure 5D; CBX mean change: -15.6%; 95% CI: 20.6% -10.6%; bootstrap resampling; 186 cells, 8 slices, 6 animals; Control mean change: +0.6%, 95% CI: - 0.7% - 2.1%; bootstrap resampling; 205 cells, 11 slices, 10 animals).

**FIGURE 5.**
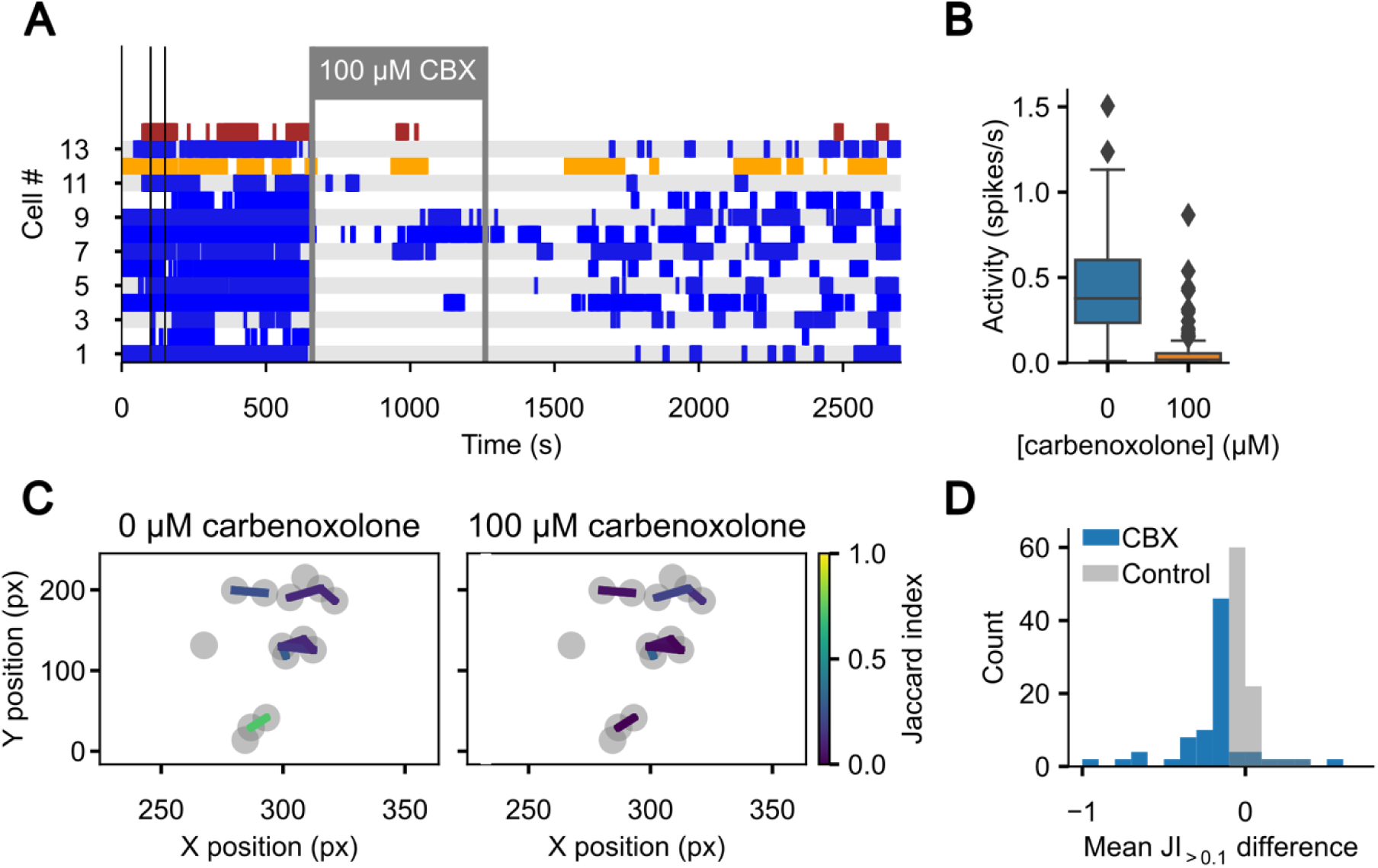
The unspecific gap junction blocker carbenoxolone reduces spiking activity, but only has minor effects on synchronized spiking. **(A)** Representative spike trains over the full recording duration of 27,000 frames (45 Minutes at 10 frames/s) of the same recording as shown in Figure 3. The enlarged section of Figure 3A is delineated by the thin vertical, black bars, and the same cells are highlighted in yellow and red. The slice was perfused with 100 µM carbenoxolone (CBX) in the extracellular solution as indicated. **(B)** Activity as determined by the number of spikes/s is reduced under perfusion with 100 µM carbenoxolone (186 cells, 8 slices, 6 animals). **(C)** Spatial distribution of the correlation between the 14 cells in (A) is shown before (left) and during perfusion with 100 µM carbenoxolone (right). Only connections with a JI above 0.1 before perfusion with carbenoxolone are shown for clarity. px – Pixels **(D)** The JI during perfusion with 100 µM carbenoxolone (CBX, blue; 186 cells, 8 slices, 6 animals) or with control solution (without CBX, grey; 205 cells, 11 slices, 10 animals) was subtracted from the JI before perfusion. A histogram of the differences for all recorded cells is shown (n(CBX): 186 cells, 8 slices, 6 animals; n(Control): 205 cells, 11 slices, 10 animals).

The strong decrease in overall spiking may be due to one of the many non-specific targets of CBX (Juszczak and Swiergiel, 2009) such as voltage-gated calcium channels (Vessey et al., 2004). The temporal correlation of spiking as indicated by the reduction JI was only marginally reduced, ruling out a large role of gap junctions in ZG calcium signaling.

## DISCUSSION

The existence of gap junctions in the ZG has been controversially discussed over the past decades (Meda et al., 1993; Usadel et al., 1993; Bell and Murray, 2016). Mutations in *CADM1* were recently associated with primary aldosteronism, and the patho-mechanism was suggested to be linked to the proper formation of connexin 43 gap junctions in the ZG (Wu et al., 2023). The functional data from that paper is mostly based on experiments in H295R cells, which have been cultured from an adrenocortical carcinoma and show some properties of all cell layers of the adrenal cortex (Bird et al., 1996). The role of gap junctions in the physiological ZG therefore remained unexplained.

We found ample evidence that connexin proteins, the building blocks of gap junctions, exist on the RNA and protein level, primarily in the form of connexin 37 and 43. We also confirmed previous reports (Guagliardo et al., 2020) of synchronized calcium oscillations among cells within ZG rosettes. This raised the hypothesis that gap junctions may – at least partially – mediate the observed correlation in activity across ZG cells. However, our functional data from calcium imaging (Figure 3) and FRAP (Figure 4B and C) only show low levels of coupling at best.

Generally, coupling between ZG cells was highest between cells in proximity (Figure 3C). These findings and the distances observed are well in agreement with previous reports that synchronization only occurs within ZG rosettes (Guagliardo et al., 2020). Coupling was previously primarily attributed to a mechanical linkage of ZG cells through beta-catenin (Guagliardo et al., 2020), which is also required for the glomerular development of cells (Leng et al., 2020). Previous studies of gap junctions using the scratch loading technique also did not show diffusion of the dye to cells across the ZG. However, this may have missed gap junctional coupling as only cells within glomerular structures may be linked, and scratch loading is therefore not a suitable method.

We performed FRAP following incubation of adrenal slice preparations with calcein AM. Recovery of the signal in a bleached cell is proportional to the density of gap junctions that allow for the passage of non-bleached dye to adjacent cells. However, these experiments only showed low levels of recovery. The degree of recovery was indistinguishable upon application of gap junction inhibitors carbenoxolone or GAP-27, indicating that the small degree of recovery observed was not due to coupling of cytosols. It may rather reflect incomplete bleaching and subsequent intracellular redistribution of the dye.

Previous studies have mainly focused on the role of Cx43 in the adrenal cortex as it is the most abundantly expressed connexin isoform in the zona fasciculata. Earlier studies also describe a lack of expression of some but not all other connexin isoforms (Bell and Murray, 2016). However, recent data from microarray studies on human (Nishimoto et al., 2015) and rat (Nishimoto et al., 2012) adrenal glands, confirmed by our data, point to significant expression of Cx37 in the ZG. We observed a distinct expression pattern for Cx37 that is highest in the blood vessels penetrating the ZG and decreases within ZG and ZF cells towards the medulla while the opposite pattern was observed for Cx43. It is intriguing to consider this in the light of the mostly centripetal development of cells from a ZG towards a ZF phenotype (Freedman et al., 2013) but comprehensive data on the functional difference between connexin isoforms and its relevance is still scarce.

Our observation that most cells showed no coupling may at least in part be due to methodological limitations. While rosettes are three-dimensional structures, calcium imaging in our study was limited to a rather thin layer and coupling to cells above or below this layer would have been undetectable. However, if the density of gap junctions were high, it would be expected that all cells within each rosette were closely coupled.

Our results cast doubt on the significance of gap junctions for ZG physiology in mice. This also leads to questions whether the pathomechanism of *CADM1* mutations (Wu et al., 2023) truly involves gap junctions. While it is possible that *CADM1* mutations upregulate the formation of gap junctions, no such evidence has been presented so far.

However, there may also be gap junction-independent roles for connexin proteins in the ZG. We observed that Cx43 in the ZG was at least partially retained intracellularly, whereas a more pronounced membrane staining was observed in the ZF (Figure 2). It has been reported that Cx43 may also directly regulate gene transcription (Dang et al., 2003; Kotini et al., 2018), so it is conceivable that this may be another role of connexins in the ZG.

## CONCLUSION

In summary, we confirmed the presence of connexin Cx43 and Cx37 in the adrenal cortex including the ZG. However, it appears that only low levels of functional gap junctions are present in the physiological murine ZG. If *CADM1* mutations indeed confer their pathological effects via gap junctions, these mutations would be expected to cause additional changes.

These may either be mediated by post-translational modifications of existing proteins or changes in gap junction gene expression. Further studies using KO models of Cx43 or Cx37 may aid in understanding their role in the ZG and the switch towards higher expression in the ZF.

## AUTHOR CONTRIBUTIONS

GS and UIS conceived the study. ISH was performed by GS and NH. IF was performed by NH and images taken by NH and GS. FRAP staining was performed by GS. GS, HAD and MV prepared and performed calcium imaging, which was analyzed by GS. The manuscript was written by GS, with revisions by all authors.

## DATA AVAILABILITY

Python analysis scripts, raw calcium signal traces, IF and ISH stainings are available on Zenodo (DOI: 10.5281/zenodo.13305189).

Raw video files are only available upon reasonable request, and transfer must be organized individually due to the large file sizes involved.

## ACKNOWLEDGEMENTS

This work was funded by the Deutsche Forschungsgemeinschaft (STO 1260/1-1 to GS, project-ID 394046635 (SFB 1365) to UIS, and project-ID 431984000 (SFB 1453) to UIS) and Stiftung Charité (BIH_PRO_406 to UIS). We thank the Advanced Medical BioImaging Core Facility of the Charité – Universitätsmedizin Berlin (AMBIO) for support in acquisition of the imaging data. We thank Sarah Döring, Nico C. Brüssow, Ana Luica Huitron Carrizales and Marie Cotta for performing mouse genotyping.

## REFERENCES

Bassett, M. H., Suzuki, T., Sasano, H., White, P. C., and Rainey, W. E. (2004). The Orphan Nuclear Receptors NURR1 and NGFIB Regulate Adrenal Aldosterone Production. Mol Endocrinol 18, 279–290. doi: 10.1210/me.2003-0005

Bell, C. L., and Murray, S. A. (2016). Adrenocortical Gap Junctions and Their Functions. Front. Endocrinol. 7. doi: 10/ghj45k

Bird, I. M., Pasquarette, M. M., Rainey, W. E., and Mason, J. I. (1996). Differential control of 17 alpha-hydroxylase and 3 beta-hydroxysteroid dehydrogenase expression in human adrenocortical H295R cells. The Journal of Clinical Endocrinology & Metabolism 81, 2171– 2178. doi: 10.1210/jcem.81.6.8964847

Capponi, A. M., Lew, P. D., Jornot, L., and Vallotton, M. B. (1984). Correlation between cytosolic free Ca2+ and aldosterone production in bovine adrenal glomerulosa cells. Evidence for a difference in the mode of action of angiotensin II and potassium. Journal of Biological Chemistry 259, 8863–8869. doi: 10.1016/S0021-9258(17)47233-4

Chaytor, A. T., Evans, W. H., and Griffith, T. M. (1998). Central role of heterocellular gap junctional communication in endothelium-dependent relaxations of rabbit arteries. The Journal of Physiology 508, 561–573. doi: 10.1111/j.1469-7793.1998.561bq.x

Christopher, G. A., Noort, R. J., and Esseltine, J. L. (2022). Connexin 43 Gene Ablation Does Not Alter Human Pluripotent Stem Cell Germ Lineage Specification. Biomolecules 12, 15. doi: 10.3390/biom12010015

Clasadonte, J., Scemes, E., Wang, Z., Boison, D., and Haydon, P. G. (2017). Connexin 43-Mediated Astroglial Metabolic Networks Contribute to the Regulation of the Sleep-Wake Cycle. Neuron 95, 1365–1380.e5. doi: 10.1016/j.neuron.2017.08.022

Dang, X., Doble, B. W., and Kardami, E. (2003). The carboxy-tail of connexin-43 localizes to the nucleus and inhibits cell growth. Mol Cell Biochem 242, 35–38.

Davidson, J. S., Baumgarten, I. M., and Harley, E. H. (1986). Reversible inhibition of intercellular junctional communication by glycyrrhetinic acid. Biochemical and Biophysical Research Communications 134, 29–36. doi: 10/cz2qgt

Davis, K. T., Prentice, N., Gay, V. L., and Murray, S. A. (2002). Gap junction proteins and cell-cell communication in the three functional zones of the adrenal gland. Journal of Endocrinology 173, 13–21. doi: 10/b84fks

Dinh, H. A., Volkert, M., Secener, A. K., Scholl, U. I., and Stölting, G. (2024). T-and L-Type Calcium Channels Maintain Calcium Oscillations in the Murine Zona Glomerulosa. Hypertension 81, 811–822. doi: 10.1161/HYPERTENSIONAHA.123.21798

Fakunding, J. L., Chow, R., and Catt, K. J. (1979). The Role of Calcium in the Stimulation of Aldosterone Production by Adrenocorticotropin, Angiotensin II, and Potassium in Isolated Glomerulosa Cells. Endocrinology 105, 327–333. doi: 10.1210/endo-105-2-327

Fern, R. J., Hahm, M. S., Lu, H. K., Liu, L. P., Gorelick, F. S., and Barrett, P. Q. (1995). Ca2+/calmodulin-dependent protein kinase II activation and regulation of adrenal glomerulosa Ca2+ signaling. American Journal of Physiology-Renal Physiology 269, F751– F760. doi: 10/ghj45w

Freedman, B. D., Kempna, P. B., Carlone, D. L., Shah, M. S., Guagliardo, N. A., Barrett, P. Q., et al. (2013). Adrenocortical Zonation Results from Lineage Conversion of Differentiated Zona Glomerulosa Cells. Developmental Cell 26, 666–673. doi: 10/f5fnvz

Friend, D. S., and Gilula, N. B. (1972). A DISTINCTIVE CELL CONTACT IN THE RAT ADRENAL CORTEX. J Cell Biol 53, 148–163.

Guagliardo, N. A., Klein, P. M., Gancayco, C. A., Lu, A., Leng, S., Makarem, R. R., et al. (2020). Angiotensin II induces coordinated calcium bursts in aldosterone-producing adrenal rosettes. Nat Commun 11, 1–15. doi: 10/ghj45b

Hu, C., Rusin, C. G., Tan, Z., Guagliardo, N. A., and Barrett, P. Q. (2012). Zona glomerulosa cells of the mouse adrenal cortex are intrinsic electrical oscillators. Journal of Clinical Investigation 122, 2046–2053. doi: 10/f32hq7

Ito, A., Ichiyanagi, N., Ikeda, Y., Hagiyama, M., Inoue, T., Kimura, K. B., et al. (2012). Adhesion molecule CADM1 contributes to gap junctional communication among pancreatic islet α-cells and prevents their excessive secretion of glucagon. Islets 4, 49–55. doi: 10.4161/isl.18675

Juszczak, G. R., and Swiergiel, A. H. (2009). Properties of gap junction blockers and their behavioural, cognitive and electrophysiological effects: Animal and human studies. Progress in Neuro-Psychopharmacology and Biological Psychiatry 33, 181–198. doi: 10.1016/j.pnpbp.2008.12.014

Kotini, M., Barriga, E. H., Leslie, J., Gentzel, M., Rauschenberger, V., Schambony, A., et al. (2018). Gap junction protein Connexin-43 is a direct transcriptional regulator of N-cadherin in vivo. Nat Commun 9, 3846. doi: 10.1038/s41467-018-06368-x

Kumar, N. M., and Gilula, N. B. (1996). The Gap Junction Communication Channel. Cell 84, 381–388. doi: 10.1016/S0092-8674(00)81282-9

Lai, S., Ma, L., E, W., Ye, F., Chen, H., Han, X., et al. (2020). Mapping a mammalian adult adrenal gland hierarchy across species by microwell-seq. Cell Regen 9, 11. doi: 10.1186/s13619-020-00042-8

Leng, S., Pignatti, E., Khetani, R. S., Shah, M. S., Xu, S., Miao, J., et al. (2020). β-Catenin and FGFR2 regulate postnatal rosette-based adrenocortical morphogenesis. Nature Communications 11, 1680. doi: 10/gg79tq

Martin, A. O., Mathieu, M.-N., Chevillard, C., and Guérineau, N. C. (2001). Gap Junctions Mediate Electrical Signaling and Ensuing Cytosolic Ca2+ Increases between Chromaffin Cells in Adrenal Slices: A Role in Catecholamine Release. J. Neurosci. 21, 5397–5405. doi: 10.1523/JNEUROSCI.21-15-05397.2001

Meda, P., Pepper, M. S., Traub, O., Willecke, K., Gros, D., Beyer, E., et al. (1993). Differential expression of gap junction connexins in endocrine and exocrine glands. Endocrinology 133, 2371–2378. doi: 10.1210/en.133.5.2371

Meme, W., Vandecasteele, M., Giaume, C., and Venance, L. (2009). Electrical coupling between hippocampal astrocytes in rat brain slices. Neuroscience Research 63, 236–243. doi: 10.1016/j.neures.2008.12.008

Munari-Silem, Y., Lebrethon, M. C., Morand, I., Rousset, B., and Saez, J. M. (1995). Gap junction-mediated cell-to-cell communication in bovine and human adrenal cells. A process whereby cells increase their responsiveness to physiological corticotropin concentrations. J Clin Invest 95, 1429–1439. doi: 10/ck77rf

Murray, S. A., Oyoyo, U. A. A., Pharrams, S. Y., Kumar, N. M., and Gilula, N. B. (1995). Characterization of GAP junction expression in the adrenal gland. Endocrine Research 21, 221–229. doi: 10/bgzfrj

Murray, S. A., and Pharrams, S. Y. (1997). Comparison of gap junction expression in the adrenal gland. Microscopy Research and Technique 36, 510–519. doi: 10/dz243w

Nishimoto, K., Rigsby, C. S., Wang, T., Mukai, K., Gomez-Sanchez, C. E., Rainey, W. E., et al. (2012). Transcriptome Analysis Reveals Differentially Expressed Transcripts in Rat Adrenal Zona Glomerulosa and Zona Fasciculata. Endocrinology 153, 1755–1763. doi: 10.1210/en.2011-1915

Nishimoto, K., Tomlins, S. A., Kuick, R., Cani, A. K., Giordano, T. J., Hovelson, D. H., et al. (2015). Aldosterone-stimulating somatic gene mutations are common in normal adrenal glands. PNAS 112, E4591–E4599. doi: 10/f7pb2p

Penton, D., Bandulik, S., Schweda, F., Haubs, S., Tauber, P., Reichold, M., et al. (2012). Task3 Potassium Channel Gene Invalidation Causes Low Renin and Salt-Sensitive Arterial Hypertension. Endocrinology 153, 4740–4748. doi: 10.1210/en.2012-1527

Pezzi, V., Clyne, C. D., Ando, S., Mathis, J. M., and Rainey, W. E. (1997). Ca2+-Regulated Expression of Aldosterone Synthase Is Mediated By Calmodulin and Calmodulin-Dependent Protein Kinases. Endocrinology 138, 835–838. doi: 10.1210/endo.138.2.5032

Schewe, J., Seidel, E., Forslund, S., Marko, L., Peters, J., Muller, D. N., et al. (2019). Elevated aldosterone and blood pressure in a mouse model of familial hyperaldosteronism with ClC-2 mutation. Nat Commun 10, 5155. doi: 10.1038/s41467-019-13033-4

Schindelin, J., Arganda-Carreras, I., Frise, E., Kaynig, V., Longair, M., Pietzsch, T., et al. (2012). Fiji: an open-source platform for biological-image analysis. Nat Methods 9, 676–682. doi: 10.1038/nmeth.2019

Seidel, E., Schewe, J., Zhang, J., Dinh, H. A., Forslund, S. K., Markó, L., et al. (2021). Enhanced Ca2+ signaling, mild primary aldosteronism, and hypertension in a familial hyperaldosteronism mouse model (Cacna1hM1560V/+). PNAS 118. doi: 10/gjsgmt

Stölting, G., Dinh, H. A., Volkert, M., Hellmig, N., Schewe, J., Hennicke, L., et al. (2023). Isradipine therapy in *Cacna1d^Ile772Met/+^* mice ameliorates primary aldosteronism and neurologic abnormalities. JCI Insight 8. doi: 10.1172/jci.insight.162468

Usadel, H., Bornstein, S. R., Ehrhart-Bornstein, M., Kreysch, H. G., and Scherbaum, W. A. (1993). Gap Junctions in the Adrenal Cortex. Horm Metab Res 25, 653–654. doi: 10/bbhzb2

Vessey, J. P., Lalonde, M. R., Mizan, H. A., Welch, N. C., Kelly, M. E. M., and Barnes, S. (2004). Carbenoxolone Inhibition of Voltage-Gated Ca Channels and Synaptic Transmission in the Retina. Journal of Neurophysiology 92, 1252–1256. doi: 10/b7rcpx

Virtanen, P., Gommers, R., Oliphant, T. E., Haberland, M., Reddy, T., Cournapeau, D., et al. (2020). SciPy 1.0: fundamental algorithms for scientific computing in Python. Nat Methods 17, 261–272. doi: 10.1038/s41592-019-0686-2

Wade, M. H., Trosko, J. E., and Schindler, M. (1986). A fluorescence photobleaching assay of gap junction-mediated communication between human cells. Science 232, 525–528. doi: 10.1126/science.3961495

Wu, X., Azizan, E. A. B., Goodchild, E., Garg, S., Hagiyama, M., Cabrera, C. P., et al. (2023). Somatic mutations of CADM1 in aldosterone-producing adenomas and gap junction-dependent regulation of aldosterone production. Nat Genet 55, 1009–1021. doi: 10.1038/s41588-023-01403-0

