## Supplementary Material for "Expression and Function of Connexin 43 and Connexin 37 in the Murine Zona Glomerulosa"

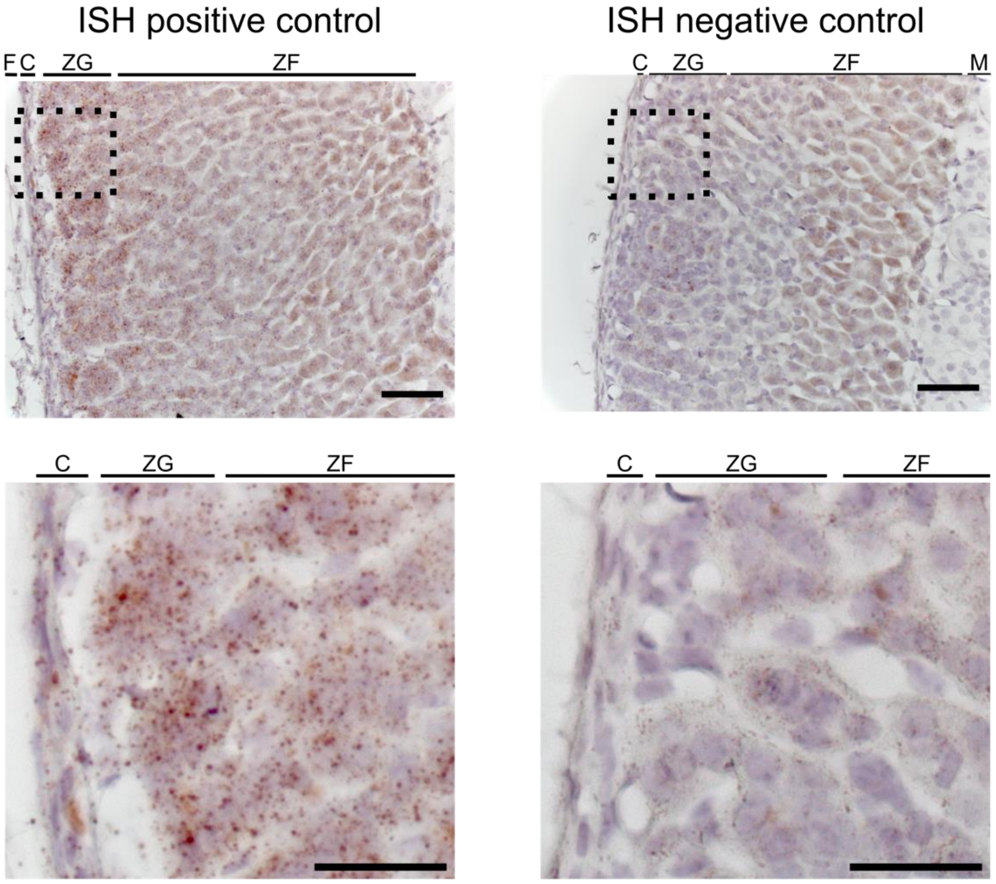


**Supplementary Figure S1.** In-situ hybridization of FFPE adrenal gland sections labeled with positive (*Ppib*) and negative control (*DapB*) probes. Lower images represent magnifications of the areas labeled with the dashed boxes in the top images. Scale bars: 50 µm (top) and 25 µm (bottom). F: fat, C: capsule, ZG: zona glomerulosa, ZF: zona fasciculata, M: medulla.

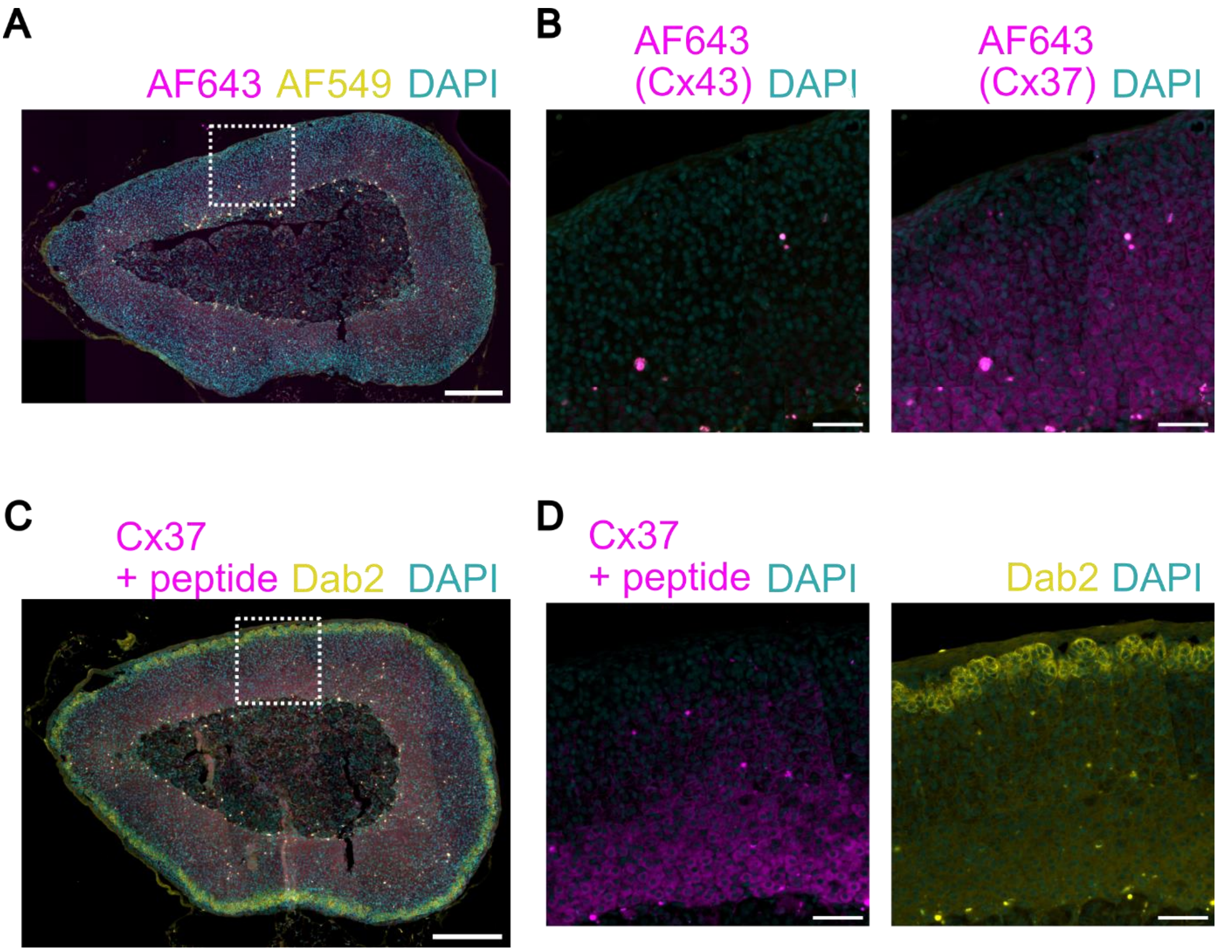


**Supplementary Figure S2.** Immunofluorescence controls. **(A)** Staining using only the secondary antibodies shows only low signals. Nuclei are stained with DAPI (cyan). The image is representative of the staining of sections (5 µm thickness) from 7 animals. Scale bar: 250 µm. **(B)** Magnification of the part enclosed by the white dashed square in A. The magenta channels have been enhanced identically to the staining of connexin 43 (left) or connexin 37 (right) as shown in Figure 2. Scale bar: 50 µm. **(C)** Immunofluorescence staining reveals the specificity of the connexin 37 staining. Preincubation with a control peptide removes staining from ZG and blood vessels as seen in Figure 2. Some signal remains in the ZF adjacent to the medulla. Nuclei are stained with DAPI (cyan). The image is representative of the staining of sections (5 µm thickness) from 8 animals. Scale bar: 250 µm. **(D)** Magnification of the part enclosed by the white dashed square in C. The magenta channel has been enhanced identically to the staining of connexin 37 as shown in Figure 2. Dab2 (yellow, right) delineates the ZG. Scale bar: 50 µm.

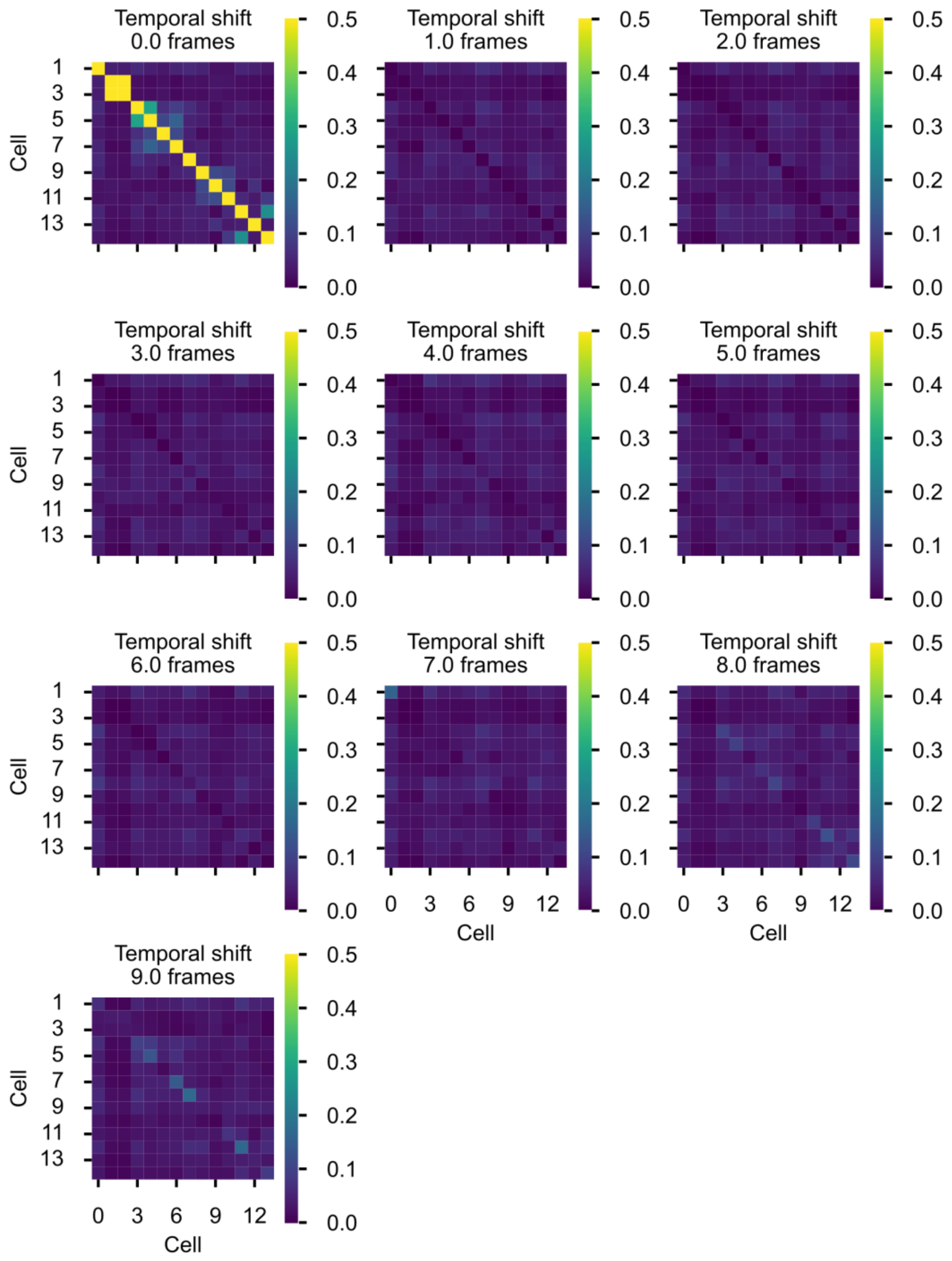


**Supplementary Figure S3.** Representative heat map of the Jaccard index values calculated between 14 cells obtained from the same recording shown in Figure 3B. JI values are color coded according to the scale shown on the right. Heat maps of the JI values were calculated between the original spike train and spike trains shifted by the indicated number of frames. No similarly high JI values as shown in the first heatmap are observed, indicating that the correlation observed is not random. Autocorrelation can be seen across the diagonal from the upper left to lower right corner.
